# A Novel Cytoskeletal Action of Xylosides

**DOI:** 10.1101/2022.01.31.478500

**Authors:** Caitlin P. Mencio, Sharada Tilve, Masato Suzuki, Kyohei Higashi, Yasuhiro Katagiri, Herbert M. Geller

**Author notes:** Schubot Center for Avian Health, Department of Veterinary Pathobiology, College of Veterinary Medicine & Biomedical Sciences, Texas A&M University, College Station, TX, United States of America.

## Abstract

Proteoglycan glycosaminoglycan (GAG) chains are attached to a serine residue in the protein through a linkage series of sugars, the first of which is xylose. Xylosides are chemicals which compete with the xylose at the enzyme xylosyl transferase to prevent the attachment of GAG chains to proteins. These compounds have been employed at concentrations in the millimolar range as tools to study the role of GAG chains in proteoglycan function. In the course of our studies with xylosides, we conducted a dose-response curve for xyloside actions on neural cells. To our surprise, we found that concentrations of xylosides in the nanomolar to micromolar range had major effects on cell morphology. These effects are due to changes in cytoskeletal dynamics. Concentrations of xylosides which were effective in altering morphology did not alter GAG chain synthesis rates, nor did they produce any changes in gene expression as determined by RNAseq of treated cells. These observations support a novel action of xylosides on neuronal cells.

## Introduction

Proteoglycans (PGs) are found in every tissue in the body. Consisting of two major components, a core protein and glycosaminoglycan (GAG) chain(s), they are essential components of the extracellular matrix. There are three major classes, differentiated by their GAG chain composition: heparan sulfate, chondroitin sulfate and keratan sulfate. PGs are involved in many important biological processes (1), especially in the central nervous system, where they regulate neuronal migration, axon guidance and differentiation (2, 3). They are also believed to play critical roles in neural de- and re-generation (4).

In the recent past, it has become more apparent that the physiological effects of proteoglycans can be attributed to their GAG chains. Moreover, the wide range of biological function is often attributed to the complexity and diversity of GAG chain modifications, sulfation being the most common. Altering GAG sulfation patterns has been known to change GAG chain receptor binding resulting in modified cell signaling (5–7).

GAG chain biosynthesis is a non-template driven process that begins for all PGs with a common linkage of three sugars (Xyl-Gal-Gal) to a serine on the core protein. The three classes then diverge with addition of disaccharides to the linkage region. For HS, these would be GlcNAc and GlcA, for CS the disaccharides are GalNAc and GlcA, and for KS they are GlcNAc and Gal. Each disaccharide in the chain may undergo several different modifications, primarily sulfation. The impact that modifications to the GAG chains has on neural development as well as other cellular processes remains an active area of research.

One major approach to understanding the function of GAG chains in PGs has been through the use of chemical modulators of GAG biosynthesis called xylosides which serve as competitive molecules for cellular sugar chain synthesis. Xyloside induced interference in GAG biosynthesis can lead to changes in GAG concentration, chain type, sulfation pattern, and molecular weight (8–10) as well as biological activity (11–13). Historically, xyloside research has utilized high (millimolar) concentrations that serve to primarily block endogenous GAG production (14, 15). Relatively few studies used lower than mM concentrations (16). We therefore conducted a dose-response study to determine the minimal concentration of xyloside that would disrupt GAG chain synthesis in hippocampal pyramidal neurons and Neuro2A cells. To our surprise, we saw major changes in cell morphology when cells were treated with nanomolar xyloside concentrations but not micro- or millimolar treatment. After documenting the cytoskeletal changes, we determined that nanomolar treatment of cells with xyoside will alter sugar profiles with distinct differences found in heparan sulfate (HS) disaccharide composition from both control cells and cells treated with the high concentration of xyloside. This study serves as a first step in our attempt to understand how minor shifts in GAG chain composition or concentration can affect biological processes and continues to support the critical role of sugars in development and cellular function.

## Materials and Methods

### Laboratory animals

Experiments and procedures were performed in accordance with Institutional Animal Care and Use Committee (IACUC) at the National Institutes of Health approved protocols. Pregnant female C57Bl/6 mice (Charles River) were housed in a pathogen free facility with standard 12 h light/dark cycle and unlimited access to food and embryonic (E17-19) pups where utilized for embryonic hippocampal neuron primary cultures.

### Cell culture

Primary hippocampal neuron cultures were prepared from embryonic (E17-19) C57Bl/6 mouse brains. Hippocampi were dissected and dissociated into single cell suspensions. Dissociated cells were seeded onto coverslips coated with poly-L-lysine and cultured in 500 μL Neurobasal medium containing B27 supplement (Thermo Fisher) and 24 mM KCl at a density of 8-10k cells/well to allow for observation of isolated neurons. After allowing 2 hr for neuronal attachment, media was replaced with 1ml fresh Neurobasal media containing DMSO, 500nM xyloside, or 1mM xyloside. Cells were incubated for 72 hr at 37°C and 5% CO2 atmosphere and then fixed and stained for DAPI, βIII-tubulin, and actin (Phalloidin).

Neuro2A (ATCC) cells were cultured in DMEM media containing 10% fetal bovine serum (FBS) at 37°C with 5% CO_2_. Cells were seeded onto coverslips, 35mm glass bottom dishes (MatTek), 6-well plates or T-75 (Corning) flasks depending on experimental design. For conditioned media, Neuro2A cells were grown to about 50% confluency in T-75 flasks (Corning) in DMEM supplemented with 10% FBS. At this point, DMEM containing FBS was removed and cells were washed thrice with sterile PBS and then kept in DMEM only overnight. The next day the media was replaced with media containing DMSO, 500nM xyloside or 1mM xyloside and cells were allowed to grow for 48h at normal cell culture conditions. At this time, media was removed and placed into 15ml conical tubes and spun for 10m at 500 rpm to remove floating cells and debris. The supernatant was transferred into a fresh tube and utilized for GAG analysis.

### Glycosaminoglycan analysis

Conditioned media was harvested from Neuro2A cells that had been serum starved overnight and then treated for 48h with xylosides in DMEM culture medium. Approximately 10ml of conditioned media was collected from each experimental condition: DMOS, 500nM and 1mM xyloside. This media was spun for 5 min at 2000 rpm to remove cell debris and then transferred to a new conical tube and frozen until analysis.

GAG extraction was performed as follows. The media (1 mL) was treated with 10 % TCA and centrifuged at 12000 rpm for 5 min to remove proteins. GAGs were collected by Amicon Ultra Centrifugal Filter 3K device (Merck Millipore, Billerica, MA, USA) and suspended with 100 μL of H_2_O. Fifty μL of GAGs solution was moved to new 1.5 mL microcentrifuge tube and lyophilized. Resulting GAG samples were incubated in the reaction mixture (35 μL) containing 28.6 mM Tris-acetate (pH 8.0) and 50 mIU of chondroitinase ABC for 16 h at 37 °C. Depolymerized samples were boiled and evaporated, unsaturated disaccharides of CS were collected by Amicon Ultra Centrifugal Filter 30K device (Merck Millipore, Billerica, MA, USA). The remaining HS samples in filters of spin columns were transferred to new microtubes and incubated in 16 μL of reaction mixture (pH 7.0), containing 1 mU heparinase I (Seikagaku Corp., Tokyo, Japan), 1 mU heparinase II (Iduron, Manchester, UK), 1 mU heparinase III (Seikagaku Corp., Tokyo, Japan), 31.3 mM sodium acetate, and 3.13 mM calcium acetate for 16 h at 37 °C. Unsaturated disaccharide analysis using reversed phase ion-pair chromatography with sensitive and specific post-column detection was performed as described previously (17). Disaccharide composition analysis of CS or HS was performed by reversed phase ion-pairing chromatography with sensitive and specific post-column detection. A gradient was applied at a flow rate of 1.0 ml min^-1^ on Senshu Pak Docosil (4.6 × 150 mm; Senshu Scientific Co., Ltd., Tokyo, Japan) at 60°C. The eluent buffers were as follows: A, 10 mM tetra-*n*-butylammonium hydrogen sulfate in 12% methanol; B, 0.2 M NaCl in buffer A. The gradient program of CS disaccharides analysis was as follows: 0-10 min (1% B), 10-11 min (1-10% B), 11-30 min (10% B), 30-35 min (10-60% B), and 35-40 min (60% B). The gradient program of HS disaccharides analysis was as follows: 0-10 min (1-4% B), 10-11 min (4-15% B), 11-20 min (15-25% B), 20-22 min (25-53% B), and 22-29 min (53% B). Aqueous (0.5% (w/v)) 2-cyanoacetamide solution and 1 M NaOH were added to the eluent at the same flow rates (0.25 ml min^-1^) by using a double plunger pump. The effluent was monitored fluorometrically (Ex., 346 nm; Em., 410 nm). Expression levels of HS or CS were expressed as total amounts of unsaturated disaccharides.

### Microscopy and Image Processing

Cells were imaged using a Nikon A1R or Zeiss 880 confocal microscope with 60X and 63X objectives depending on experiment. Z-stacks were maximally projected onto a single plane using Zeiss or ImageJ image processing software. Fixed cell imaging: For images used in fluorescence quantification, image capture settings were held constant, and samples from within each group were imaged at the same time. Fluorescence intensity was measured using ImageJ, with identical settings for all samples within each analysis.

### Neurite outgrowth and growth cone analysis

After fixation and staining, at least 60 images were taken across two coverslips per condition. Files were analyzed by an experimenter blinded to the experimental conditions. Neurons were measured if they were isolated from other neurons and had distinct nuclei and at least one neurite longer than the diameter of the cell body. Both longest and total neurite measurements were obtained for each neuron and at least 60 neurons were measured for each condition. Growth cone measurements were conducted by tracing the end of the neurite from the point it widened from its average diameter of the stalk and where phalloidin staining began to show more intense and spiked appearance as is common with growing ends of neurites. Only the largest growth cone of the neuron was measured. Each experiment was performed in triplicate.

### Analysis of cytoskeleton dynamics

EB3 comet analysis: Neuro2A cells were grown in culture until about 30% confluent. Cells were then treated with DMSO, 500nM xyloside or 1mM xyloside for 48h. 24h into the incubation, cells were transfected with EB3-GFP using Avalanche^®^-Omni transfection reagent. The next day, cells were imaged on the Nikon A1R with a 60x oil objective. Transfected cells were imaged every12.5 sec over 5 min. Images were then processed using u-track software (18) and comets assessed for speed, lifetime and length.

Actin bundle analysis: Neuro2A cells were grown in culture until about 30% confluent. Cells were then treated with DMSO, 500nM xyloside or 1mM xyloside for 48h. 24h into the incubation, cells were transfected with Ftractin-mCherry using Avalanche^®^-Omni transfection reagent. The next day, cells were imaged on the Zeiss 880 confocal microscope with 63X objectives. Transfected cells with lamellipodia were imaged in a z-stack. In FIJI, z-stacks were max projected and a line drawn through the lamellipodia and a line scan performed based upon fluorescence. Actin bundles were identified as peaks of increased fluorescence. Bundles were counted and the area under the curve taken to compare quantity and size of the bundles.

### RNA-seq

Between 6-15 micrograms per sample of total RNA from three samples each of Neuro2A cells treated with either DMSO, LCX or HCX were sequenced by Illumina at Omega Bioservices (Norcross, GA). Data analysis was performed using the Partek Flow statistical analysis software (Partek Incorporated). For this, raw data from three replicates of the three conditions were imported to Partek Flow for alignment and quality controls. Aligned reads were quantified to transcriptome, filtered out on low expression and normalized. Feature gene lists with at least a two-fold change in gene expression with a P < 0.05 were created by pairwise comparison. Pseudogenes were eliminated from the final list of altered genes and the list plotted as a heatmap. In order to visualize the difference of the expression between DMSO, LCX and HCX, the data were centered by subtracting the mean of the log2 Fold Change of all samples for each gene from the original log2 Fold Change value. The centered data of all samples were then plotted into a heat map The rows of the heatmaps (genes) were ordered by foldchange (19).

### Statistics

All statistical tests were performed using GraphPad Prism 7.0 (GraphPad Software, La Jolla, CA). Neurite lengths in culture were compared using Kruskal-Wallis Analysis of Variance and Mann-Whitney U tests.

## Results

We sought to evaluate the effects of inhibition of GAG chain synthesis on hippocampal neurons in culture. A previous study noted that concentrations of 0.1 and 0.2 mM xyloside perturbed the generation of neuronal polarity (20), but we sought to determine a more complete dose-response curve. We therefore added 4-methyl-umbelliferyl-β-D-xylopyranoside (4-MU) in a range of 0.5 nM to 1mM to the medium of dissociated embryonic hippocampal neurons, and fixed and stained cultured after 72h. We observed a surprising dose-dependent response to 4-MU: neurons treated with lower concentrations showed enlarged growth cones that exhibited looped microtubules as compared to DMSO treated cultures or cultures treated with mM concentrations (Fig. 1, Supp. Fig. 1). Fig. 1A shows representative images taken from the range of concentrations: DMSO (control), 500 nM 4-MU as the low xyloside concentration (LCX) and 1 mM MU as the high xyloside concentration (HCX). As presented in Fig. 1B, there was a difference in growth cone size with the different treatments: neurons treated with LCX had significantly larger growth cones than those treated with DMSO or HCX. To determine if the enlarged growth cones was due to neurite stalling, the length of the longest neurite was measured for each neuron in the condition. There was no significant difference in neurite length between all conditions (Fig. 1C). The lower end of the dose-response curve is presented in Supp. Fig. 1.

**Figure 1.**
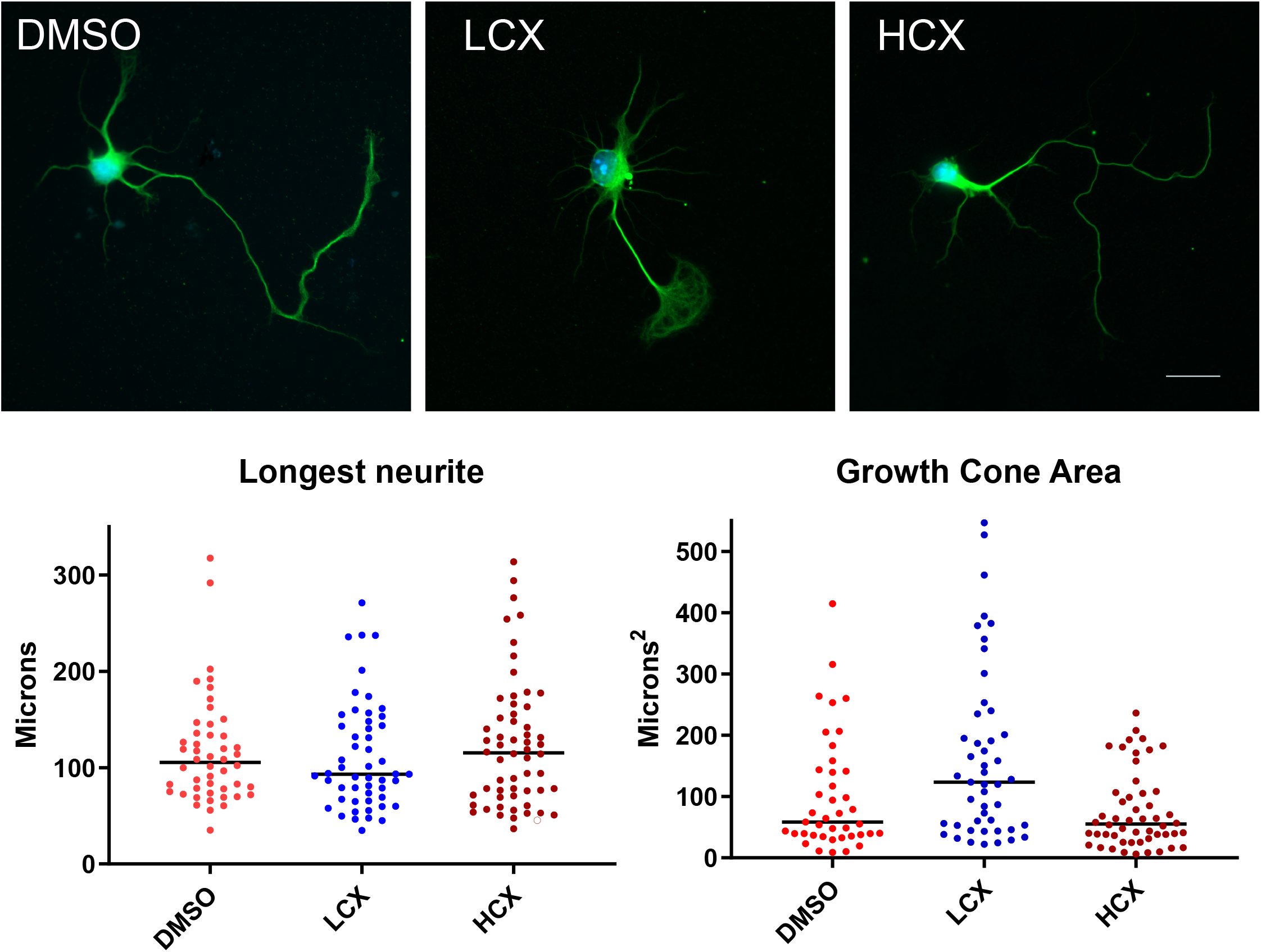
Altered cytoskeleton seen in cells treated with 500nM (LCX) but not 1mM (HCX) xyloside. A) Example of a primary mouse hippocampal neuron treated for 72h with DMSO, LCX, or HCX. White arrow indicates enlarged growth cones in LCX treated neuron. B) Quantification of growth cone area and length of longest neurite. in DMSO, LCX and HCX treated neurons. LCX treated neurons have significantly increased growth cone area compared to control or HCX conditions. Scale = 25 μm. \*\**p* < 0.01

As primary neuron culture is often limiting in cell number, we decided to assess if the mouse neural crest-derived cell line Neuro2a (N2A) would show similar morphological changes and could be used to study any structural or biochemical changes between HCX and LCX treatment. Observation of the cytoskeleton and overall cellular morphology in LCX treated N2A cells shows that these cells exhibit actin-rich lamellipodia that were not observed in N2A cells treated with either DMSO or HCX (Supp. Fig. 2). With a confirmed change in cytoskeleton and for the sake of uniformity and economy, we decided to utilize N2A cells for all other experiments in this study.

### LCX treatment reduces cellular movement

After observing changes in morphology, we next wanted to check if these changes alter the cells ability to migrate. N2A cells were treated with DMSO, LCX or HCX for 48 h at which point cells were imaged using brightfield microscopy for 4h with images being taken every 8-10 min (Fig. 2). Cells within the image were tracked and velocity and total distance was calculated. There was a significant reduction in both velocity and total distance in LCX treated cells as compared to DMSO and HCX treated N2As (Fig. 2B). This reduction in movement implicates possible changes in cytoskeleton dynamics.

**Fig. 2.**
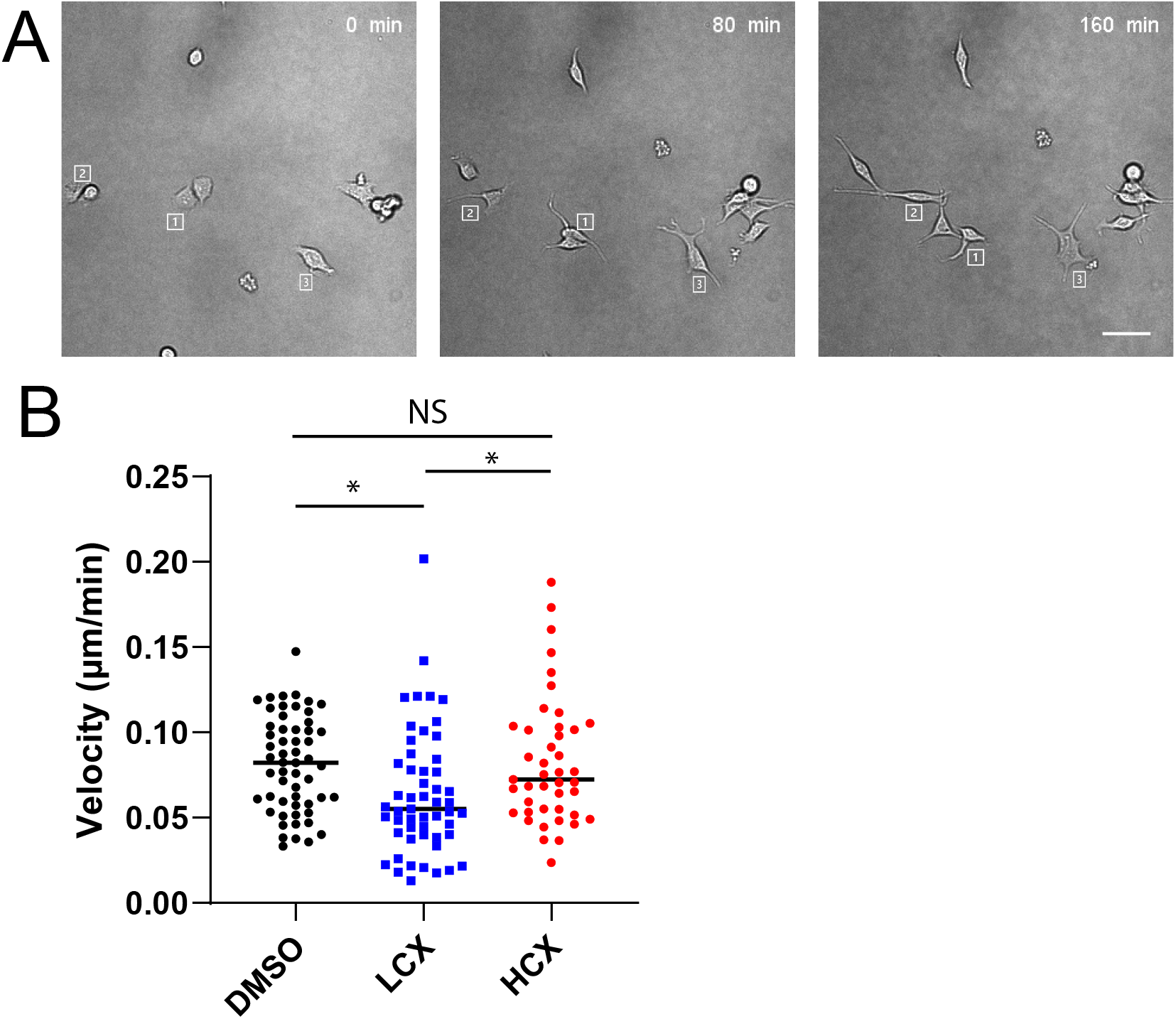
LCX reduces cell migration. Neuro2A cells were observed using time-lapse microscopy. A) Representative images taken at the indicated intervals. Letters show positions of specific cells. B) Dot plot of velocity of individual cells taken as distance moved in each frame. \**p* < 0.05. Scale = 25 μm

### LCX treatment affects both actin and microtubule dynamics

To determine if altered cytoskeleton may plan a role in LCX induced changes in morphology and migration, we next assessed both actin and microtubules in N2A cells. N2A cells were transfected with Ftractin-mCherry and treated for 48h with DMSO, LCX or HCX. At 48h, visual observation showed a marked difference in lamellipodia of LCX treated cells as compared to DMSO and HCX treated N2As. DMSO and HCX-treated cells showed bright and thick actin bundles while LCX treated cells appeared to have fewer bundles which also appeared thinner than their control counterparts (Fig. 3A). We quantified both the area and number of actin bundles from these images. LCX treated N2A cells exhibited lamellipodia that had significantly fewer actin bundles per 10μm when compared to controls (Fig. 3B). Additionally, the area under the curve for these bundles was significantly reduced in LCX treated cells as compared to DMSO or HCX treated N2As (Figure 2C) indicating less robust actin bundles in LCX lamellipodia.

**Figure 3.**
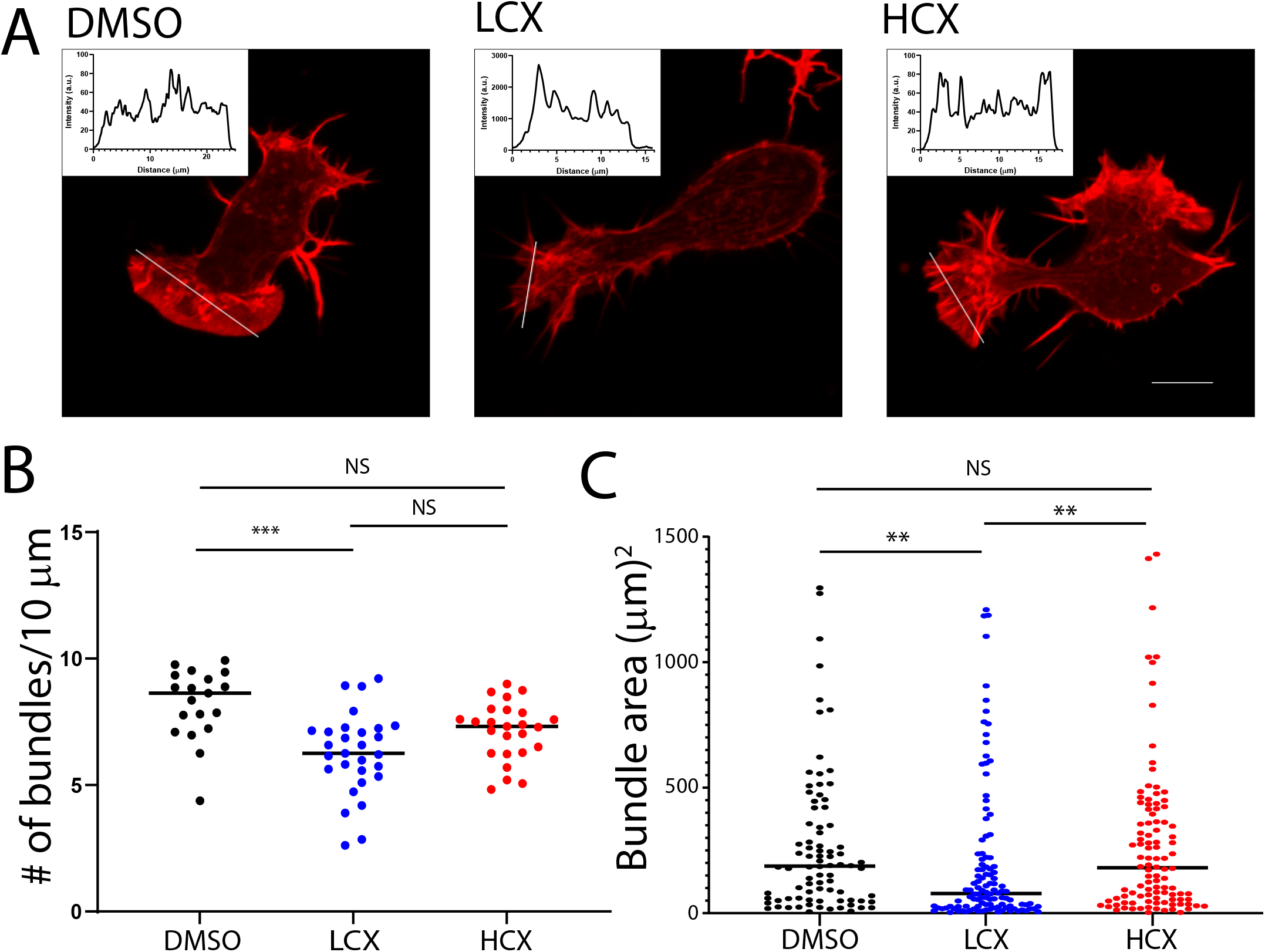
LCX alters actin bundling. A) Representative images of Neuro2A cells treated with DMSO, LCX, or HCX, and fixed and stained with phalloidin. Insets show line scans of fluorescence intensity across the lamellipodia of indicated cells. B) Plots of area under the curve and the number of peaks for each of the three conditions. Scale = 10 μm. \*\**p* < 0.01.

With a measured effect on actin, we next wanted to assess if microtubules, the other major cytoskeleton element, were also affected. To examine microtubule dynamics, EB3-GFP transfected N2A cells were treated for 48h with DMSO, LCX or HCX. EB3-GFP cells were imaged at 60x with one image taken every 12.5 sec for 5 min (Fig. 4A). Images were used to measure speed, persistence and lifetime. Both LCX and HCX treated N2A cells exhibited a higher percentage of faster moving EB3 comets as compared to controls (Fig. 4B, left). Additionally, xyloside treatment resulted in longer persistence of EB3 comets compared to DMSO treated N2As (Fig. 4B, center). There were no significant changes in comet lifetime between all conditions (Fig. 4B, right). These results, taken together with the effects on actin bundling, suggest LCX treatment alters cytoskeleton dynamics which lead to changes in cellular morphology and movement.

**Figure 4.**
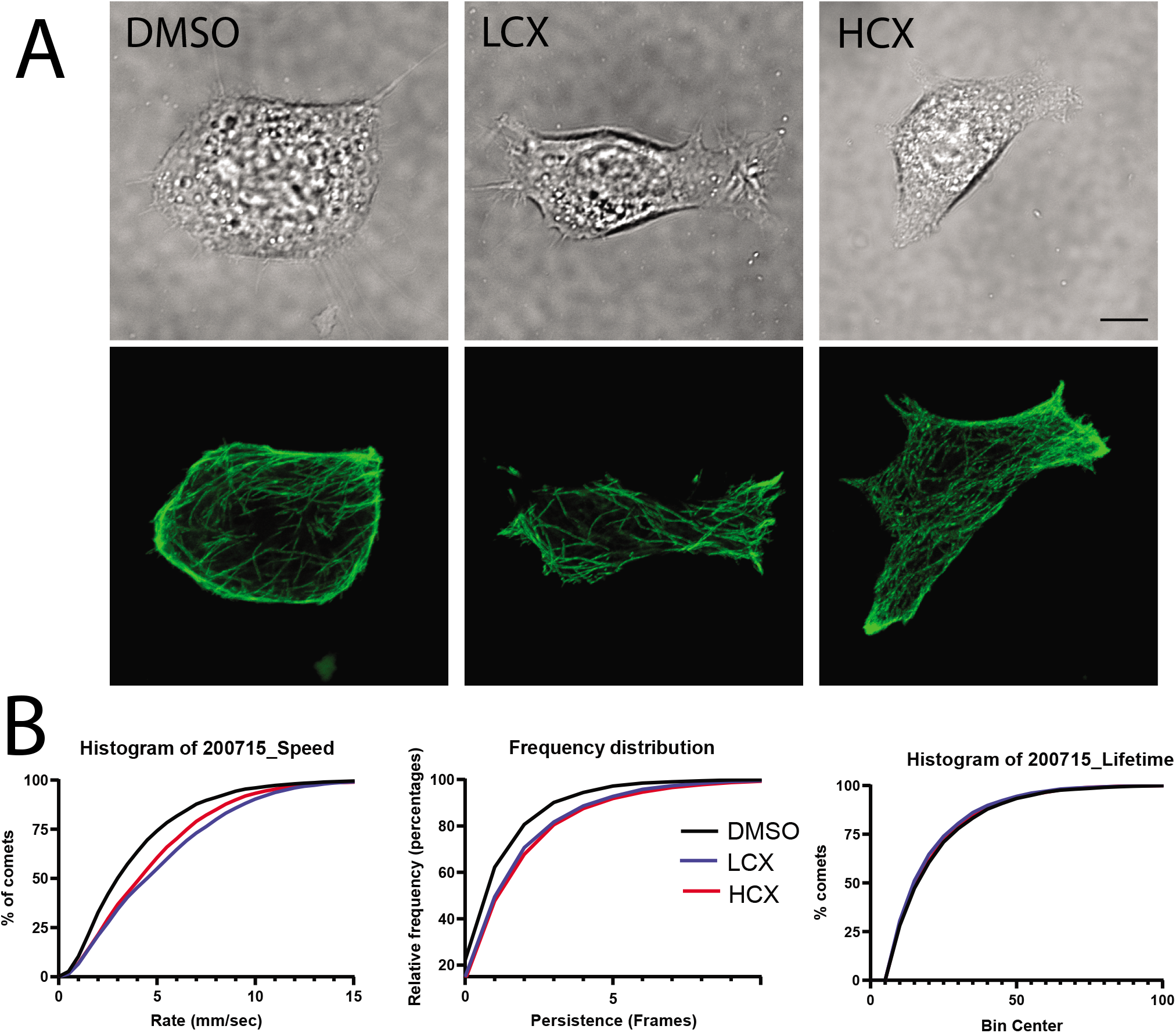
Xyloside treatment alters microtubule dynamics. A) Images of living Neuro2A cells (top) expressing EB3-EGFP were imaged over time. Bottom images show z-projections of 25 frames of EB3-EGFP. B) Cumulative distribution plots of speed, persistence and comets from DMSO, LCX and HCX-treated cells. Scale = 10 μm

### LCX treatment alters GAG chain but not mRNA profile

Xylosides have been known to affect cellular GAG chains, both increasing the rate of synthesis and secretion (21) and composition (22). These GAG chains have been linked to cellular signaling that could lead to altered cytoskeleton. To determine if LCX treatment affects GAG chain production or sulfation, N2A cells were treated with DMSO, LCX, or HCX in serum free media for 48h. Conditioned media were collected and GAG chains analyzed for concentration and disaccharide composition. As previously reported, there was a dose dependent increase in GAG chain accumulation with xyloside treatment. Both LCX and HCX increased CS chain concentration as compared to control. Notably, HCX caused a much larger increase that was significantly different from LCX (Fig. 5A).

**Figure 5.**
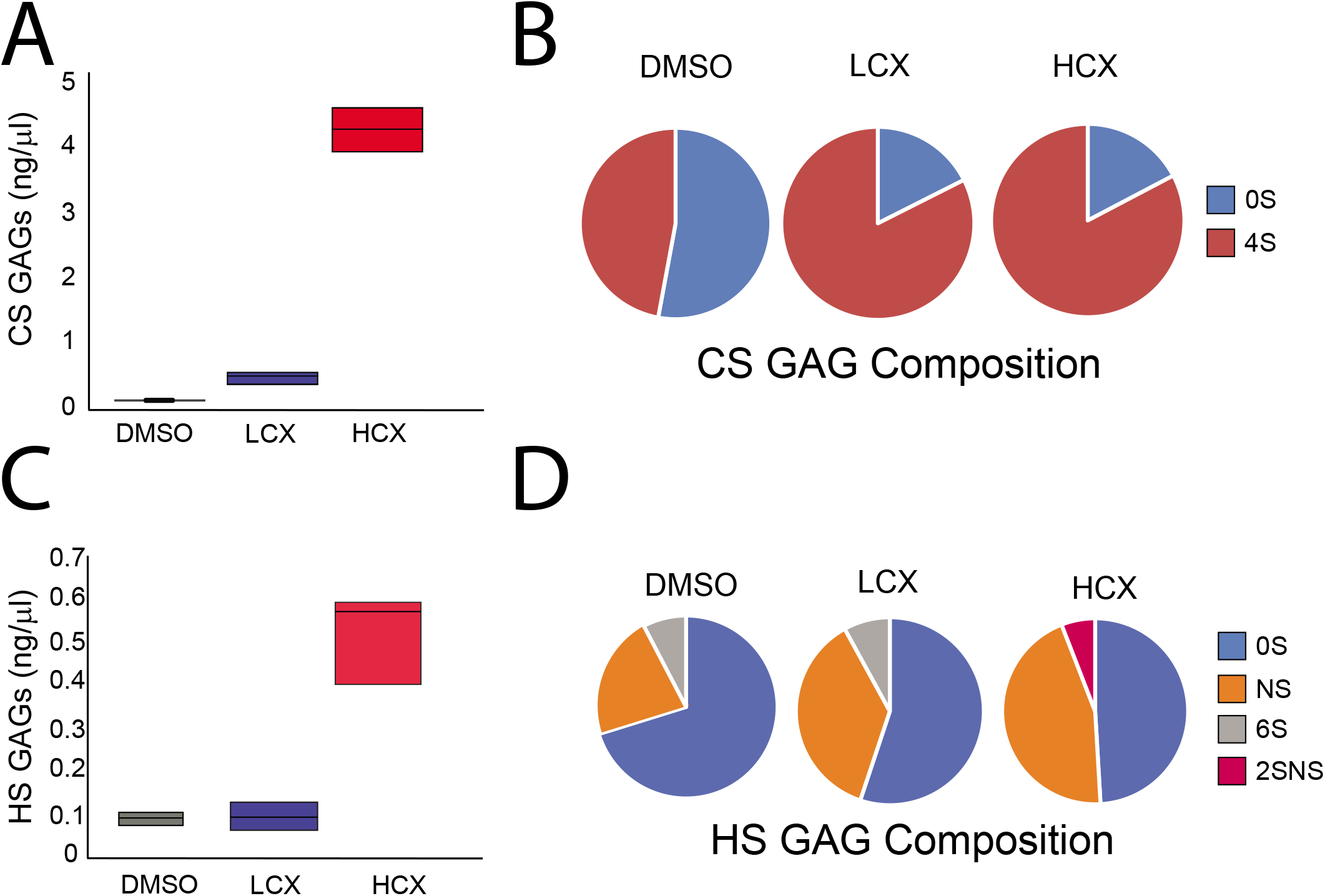
Analysis of glycosaminoglycan chains in conditioned media from Neuro2A cells after xyloside treatment. A) Quantification of CS GAGs present in conditioned media sample. B) CS disaccharides analysis C) Quantification of HS GAGs present in conditioned media sample. D) HS disaccharide analysis.

Disaccharide composition showed that conditioned media from DMSO treated cells contained a close to even split between non-sulfated and 4-sulfated chondroitin sulfate GAG chains. Xyloside treatment, both LCX and HCX, lead to an increase in 4-sulfated CS GAG (Fig, 5B). However, both LCX and HCX showed the same change while the changes in morphology and cytoskeleton were quite different. This could be due to changes in HS GAG chains, the next most abundant sugar chain that is often involved in receptor binding.

The HS chains found in the conditioned media were also assessed. In contrast to CS, only HCX treatment led to a significant increase in HS as compared to DMSO and LCX. There was no difference between HS concentration of DMSO and LCX (Fig. 5C). However, upon examination of the disaccharide composition, it was found that all three treatment groups had different disaccharide profiles. Conditioned media from DMSO treated cells showed mostly non-sulfated disaccharides with a small percentage of N-sulfated and even smaller group of 6-sulfated disaccharides (Fig. 5D). In contrast, LCX treatment resulted in an increase in N-sulfated disaccharides while HCX treatment led to a larger increase in N-sulfation, loss of 6-sulfation and the presence of a small amount of 2-sulfated, N-sulfated disaccharides (Fig. 4D). These findings indicate that xyloside treatment can change GAG chain synthesis and composition, and alterations to HS appear to occur in a concentration dependent manner.

To rule out the possibility that changes in cytoskeleton observed may be due to LCX treatment altering transcription and thus protein levels, we next performed RNA-seq to determine which, if any, proteins displayed altered mRNA production. Cultures of Neuro2A cells were treated with DMSO, LCX and HCX and RNA was extracted after 48 hours and pyrosequenced. The raw data were filtered using Partek Flow as noted in materials and methods. The filtered counts were then used to determine those genes whose expression was altered by >2X and statistically significant using p<0.05 FDR. Using these criteria, we found no genes whose change in expression was different between LCX and DMSO, while we did find changes in gene expression when comparing HCX with both DMSO and LCX. Fig. 6A presents volcano plots which indicate the differentially expressed genes between HCX and DMSO and HCX and LCX. Fig. 6B presents a Venn diagram that indicates the number of genes changed in each condition. Interestingly, majority of genes whose expression was changed were downregulated by exposure to xylosides (Fig. 6B). The Venn diagram also indicates that only about 1/3 of the genes were differentially expressed by both HCX vs DMSO and HCX vs LCX. Fig. 6C presents a heat map of the level of expression of those genes which are known to be translated into protein as compared to the average level over all conditions. There was no preference in in specific pathways using GO terms. Thus, the changes in cytoskeletal dynamics due to LCX cannot be due to changes in gene expression.

**Figure 6.**
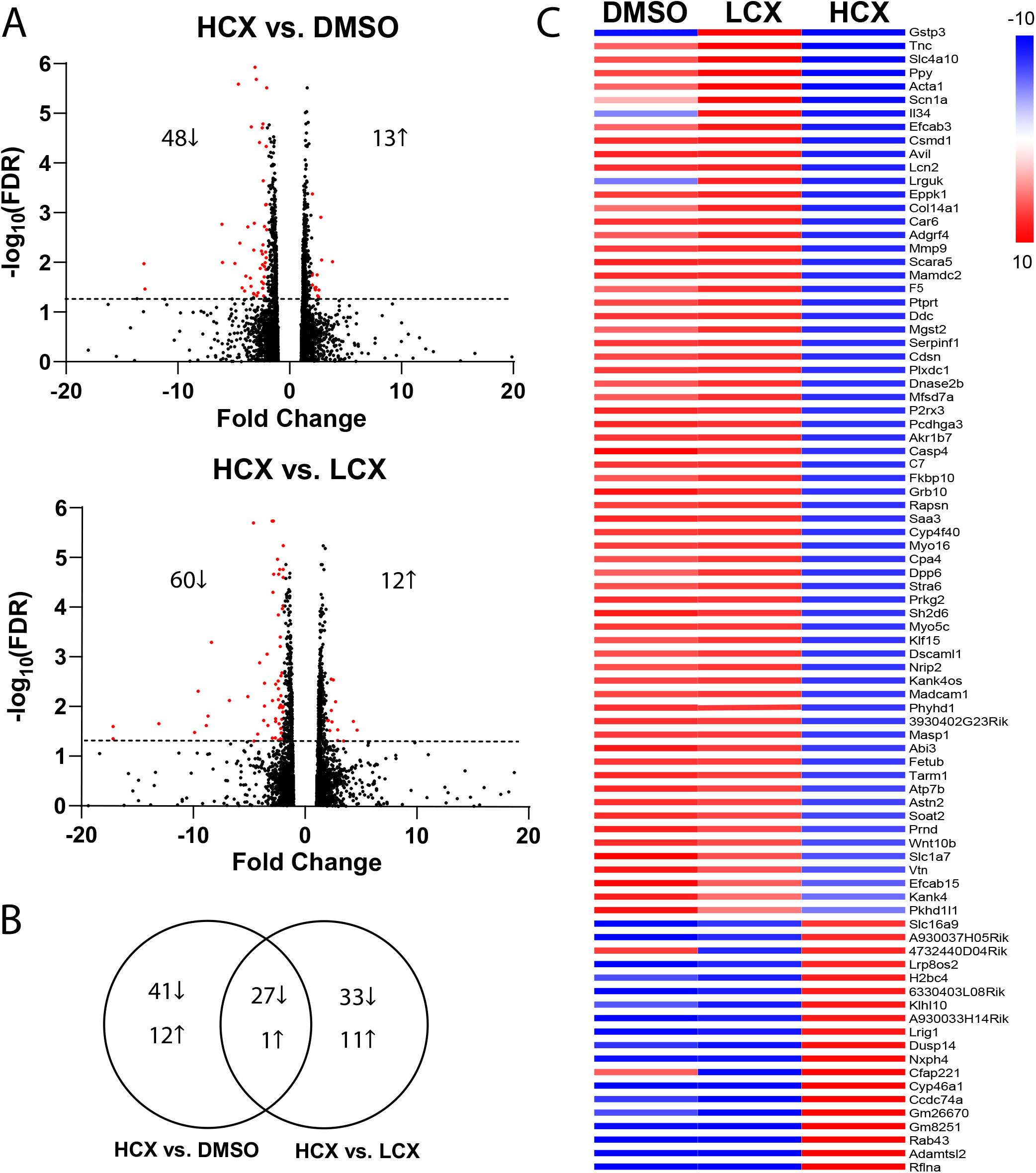
RNAseq analysis of xyloside-treated Neuro2A cells. Neuro2A cells were treated with the indicated levels of xylosides for 72h. RNA was extracted and sequenced. A) Volcano plots showing significant changes in gene expression (>50% change, P < 0.5) between HCX and DMSO-treated cells and HCX and LCX-treated cells. B) Venn diagram showing numbers of genes changed with HCX compared to DMSO and LCX treatment. C) Heat map of expressed genes that were changed by xyloside treatment. Color represents deviation from the average expression over all conditions.

## Discussion

Xylosides have been used in research as a GAG-biosynthesis inhibitor since the 1970s (14), and continue to be used today. Virtually all published work has used these small molecules in the millimolar range as originally published by Schwartz et al. (21). Our work reveals a novel action of xylosides on cell morphology that occurs at sub-micromolar concentrations, well below any previously reported to be active. Interestingly, LCX treatment only slightly raised the secretion of CS and HS into the medium, a major effect of these compounds when used in culture. Moreover, we could not identify any changes in gene expression in cells exposed to 4-MU.

Several studies have investigated dose-response curves with xylosides GAG chain production on at concentrations in the micromolar range. Carrino and Caplan (10) found increased incorporation of ^35^S into the medium of chick muscle cultures with concentrations down to 1 C. Interestingly, this concentration had minimal effect on the structure of CS GAG chains, while mM concentrations significantly altered structure. Similarly, Weinstein, et al. (23) found increased ^35^S secretion into the medium by human chondrocytes treated with 25 μM xylosides, with a maximum attained at 100 μM. More recently, Persson, et al. (22), systematically looked at GAG chain synthesis and composition influenced by (D)2-naphthyl β-D-xylopyranoside (XylNap) in several different cell lines. Consistent with other observations, they found an increase in GAG production with 10 μM XylNap, peaking at 100 μM. They also found both concentration and cell-type changes in HS and CS. These results are consistent with our findings that treatment with concentrations of 4-MU lower than 1 μM did not alter GAG chain production.

The major change we found with low concentration of 4-MU were in the cytoskeleton. Several previous studies have examined the effect of high concentrations of xylosides on cytoskeleton. Adding 1mM xyloside to vascular smooth muscle cells lead to a reduction in the number of α-actin containing cytoskeletal filaments (24, 25). Treatment with 2mM xyloside treatment impeded tubule formation during nematocyst development, implicating CS in the stabilization of membrane protrusions (26). Several studies have found that inhibiting GAG biosynthesis often leads to a reduction in invasion and migration, implicating the extracellular matrix as a modulator for cytoskeletal organization or stability (27–29) In one of the few research studies that utilized lower concentrations of xyloside, Mani et al. (30) showed that treatment with micromolar xyloside inhibited cell proliferation, a cytoskeletal dependent process, in a range of cell types. They also showed that this effect was cell type specific with 150-200 μM xyloside resulting in 50% inhibition of proliferation in human lung carcinoma, 50 μM xyloside treatment in transformed human umbilical vein endothelial cells, and no treatment effect in human lung fibroblasts. Because these manipulations likely altered both HS and CS GAGs, the question does arise as to whether these changes were due to alterations in growth factor signaling.

Previous research in our lab as well as others has shown that neurons and neuronal cell types are sensitive to GAG chains and changes in sulfation patterns (31–33). This sensitivity is linked to several processes that are regulated by cytoskeletal rearrangements such as neural migration, polarity and axon guidance. For example, degradation of GAG chains by enzyme or disruption of biosynthesis by xylosides leads to altered neuronal migration (34). The addition of HS and CS to embryonic rat neurons primarily resulted in neurons with a single long axon. Conversely the addition of DS resulted in neurons with increased dendritic growth that maintained higher levels of microtubule-associated protein 2 expression (35). In terms of axon guidance, many studies have shown the bifunctionality of HS and CS as guidance cues. HS is commonly associated with permissive substrates while CS, especially 4-sulfated CS, is seen as inhibitory (36). Proteins that have been implicated in axon guidance such as Sema5A and RPTPσ have been shown to interact with both CS and HS at binding sites unique to the type of GAG chain (7, 37). As both Sema5A and RPTPσ have been shown to influence cytoskeletal signaling pathways, it would not be a leap to associate GAG binding with changes in the neural cytoskeleton. The changes in morphology we observed would support the idea that altering GAGs could result in aberrant signaling which may affect neuronal cytoskeleton. Additionally, our results suggest that the difference in changes present between low and high concentration xyloside treatment may lean toward subtle changes in GAG chains having more distinct biological effects. This is an area of research that needs further exploration.

Because the morphological changes in neurons take several days to develop, we tested the hypothesis that there would be changes in gene expression. However, our RNAseq analysis did not find any RNAs whose level was significantly changed by exposure to LCX as compared to the control situation. However, we did find a subset of genes whose expression was significantly increased by HCX over either control or LCX. A large group of these code for proteins in the extracellular matrix, including Tenascin-C, Collagen 14, MMP9 and astrotactin 2, while another group, including a-actin, advillin, protein tyrosine phosphatase receptor type T and ABI family member 3 are involved in controlling cytoskeletal dynamics. However, none of these were changed at concentrations that produced the morphological changes. Treatment of HeLa cells with mM xylosides caused the upregulation of several RNAs related to proteoglycan synthesis (38), but we did not find any overlap with our results. Thus, it is not likely that the changes we observe are due to changes in gene expression.

As low concentration xyloside treatment alters concentration and sulfation patterns of both HS and CS GAG chains, it is possible that observed effects could be driven by multiple pathways. These changes appear to be concentration specific. No visible effect is observed in 1mM xyloside treated neurons or neuro2A cells. This could imply that utilizing lower concentrations that result in distinct, but subtle changes to GAG profiles may produce more unique and useful biological outcomes. By not overwhelming the biosynthetic machinery, nanomolar concentrations may provide differing GAG chains that can work with still present endogenous GAG chains to support or inhibit GAG mediated receptor binding. As low concentration xyloside treatment does not appear to alter transcription, we believe the next steps are to focus on assessing any phosphorylation changes in signaling pathways linked to cytoskeletal control. Further research is needed to determine the effects of LCX on intracellular signaling pathways controlling cytoskeletal rearrangements.

## Acknowledgements

Images in this manuscript were acquired in the Light Microscopy Core of the Division of Intramural Research of the National Heart, Lung, and Blood Institute, NIH. We greatly appreciate advice and assistance from Dr. Hiro Katagiri.

---

**Supp. Figure 1.**
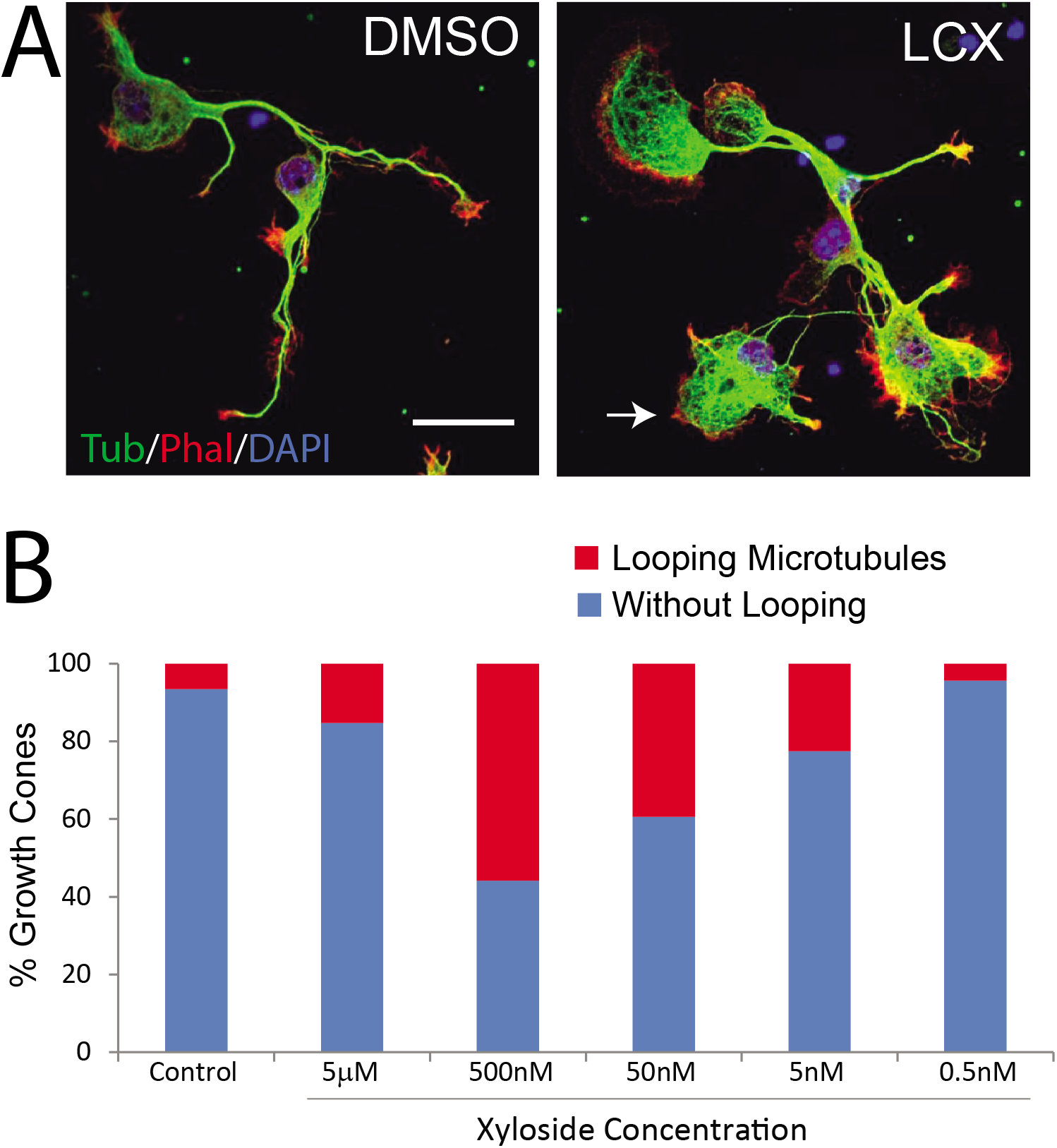
Microtubule looping in xyloside-treated growth cones. A) Images of hippocampal neurons treated with either DMSO or LCX. LCX-treated neurons have large growth cones with extensive microtubule looping (arrow). B) Dose response curve for xyloside treatment. Percentage of neurons with looped microtubules at the end of the growth cones increased and peaked at 500nM xyloside treatment.

**Supp. Fig. 2.**
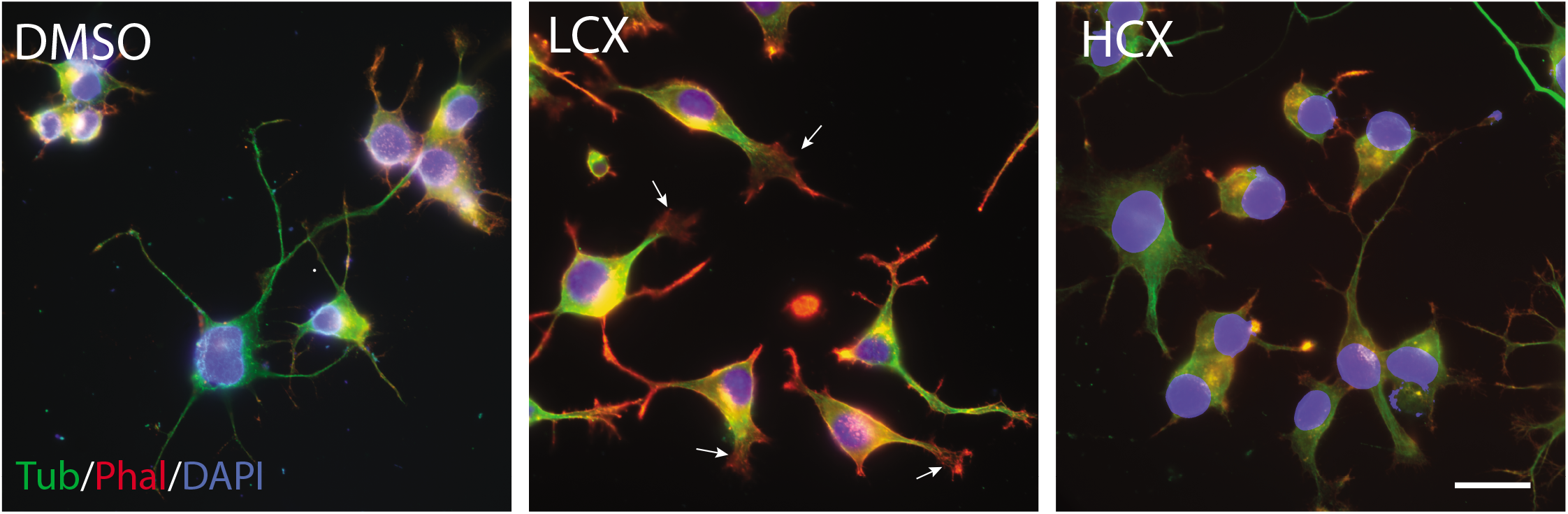
Altered Neuro2A morphology in cells treated with LCX or HCX. (Left) DMSO-treated Neuro2A cells show typical morphology with extended processes. (Center) LCX-treated cells show increased levels of phalloidin staining with lamellipodia (arrow). (Right) HCX-treated cells resemble DMSO-treated cells with processes. Scale = 25 μm.

## References

1. Wight TN, Kinsella MG, Qwarnstrom EE. The role of proteoglycans in cell adhesion, migration and proliferation. Curr Opin Cell Biol. 1992;4(5):793–801.

2. Maeda N. Proteoglycans and neuronal migration in the cerebral cortex during development and disease. Front Neurosci. 2015;9:98.

3. Masu M. Proteoglycans and axon guidance: a new relationship between old partners. J Neurochem. 2016;139 Suppl 2:58–75.

4. Bandtlow CE, Zimmermann DR. Proteoglycans in the developing brain: new conceptual insights for old proteins. Physiol Rev. 2000;80(4):1267–90.

5. Salbach J, Kliemt S, Rauner M, Rachner TD, Goettsch C, Kalkhof S, et al. The effect of the degree of sulfation of glycosaminoglycans on osteoclast function and signaling pathways. Biomaterials. 2012;33(33):8418–29.

6. Miller GM, Hsieh-Wilson LC. Sugar-dependent modulation of neuronal development, regeneration, and plasticity by chondroitin sulfate proteoglycans. Exp Neurol. 2015;274(Pt B):115–25.

7. Katagiri Y, Morgan AA, Yu P, Bangayan NJ, Junka R, Geller HM. Identification of novel binding sites for heparin in receptor protein-tyrosine phosphatase (RPTPσ): Implications for proteoglycan signaling. J Biol Chem. 2018;293(29):11639–47.

8. Schwartz NB, Domowicz MS. Proteoglycans in brain development and pathogenesis. FEBS Lett. 2018;592(23):3791–805.

9. Fritz TA, Esko JD. Xyloside priming of glycosaminoglycan biosynthesis and inhibition of proteoglycan assembly. Methods Mol Biol. 2001;171:317–23.

10. Carrino DA, Caplan AI. The effects of β-D-xyloside on the synthesis of proteoglycans by skeletal muscle: lack of effect on decorin and differential polymerization of core protein-bound and xyloside-linked chondroitin sulfate. Matrix Biol. 1994;14(2):121–33.

11. Smith-Thomas LC, Stevens J, Fok-Seang J, Faissner A, Rogers JH, Fawcett JW. Increased axon regeneration in astrocytes grown in the presence of proteoglycan synthesis inhibitors. J Cell Sci. 1995;108(Pt 3):1307–15.

12. Muto J, Naidu NN, Yamasaki K, Pineau N, Breton L, Gallo RL. Exogenous addition of a C-xylopyranoside derivative stimulates keratinocyte dermatan sulfate synthesis and promotes migration. PLoS One. 2011;6(10):e25480.

13. Raman K, Ninomiya M, Nguyen TK, Tsuzuki Y, Koketsu M, Kuberan B. Novel glycosaminoglycan biosynthetic inhibitors affect tumor-associated angiogenesis. Biochem Biophys Res Commun. 2011;404(1):86–9.

14. Schwartz NB. Regulation of chondroitin sulfate synthesis. Effect of β-xylosides on synthesis of chondroitin sulfate proteoglycan, chondroitin sulfate chains, and core protein. J Biol Chem. 1977;252(18):6316–21.

15. Okayama M, Kimata K, Suzuki S. The influence of *p*-nitrophenyl ß-d-xyloside on the synthesis of proteochondroitin sulfate by slices of embryonic chick cartilage. J Biochem. 1973;74(5):1069–73.

16. Schwarz K, Breuer B, Kresse H. Biosynthesis and properties of a further member of the small chondroitin/dermatan sulfate proteoglycan family. J Biol Chem. 1990;265(35):22023–8.

17. Imamura M, Higashi K, Yamaguchi K, Asakura K, Furihata T, Terui Y, et al. Polyamines release the let-7b-mediated suppression of initiation codon recognition during the protein synthesis of EXT2. Sci Rep. 2016;6:33549.

18. Roudot P, Legant WR, Zou Q, Dean KM, Welf ES, David AF, et al. u-track 3D: measuring and interrogating intracellular dynamics in three dimensions. bioRxiv. 2020:2020.11.30.404814.

19. Zhu G, Mayer-Wagner S, Schroder C, Woiczinski M, Blum H, Lavagi I, et al. Comparing effects of perfusion and hydrostatic pressure on gene profiles of human chondrocyte. J Biotechnol. 2015;210:59–65.

20. Nishimura K, Ishii M, Kuraoka M, Kamimura K, Maeda N. Opposing functions of chondroitin sulfate and heparan sulfate during early neuronal polarization. Neuroscience. 2010;169(4):1535–47.

21. Schwartz NB, Galligani L, Ho PL, Dorfman A. Stimulation of synthesis of free chondroitin sulfate chains by ß-D-xylosides in cultured cells. Proc Natl Acad Sci USA. 1974;71(10):4047–51.

22. Persson A, Ellervik U, Mani K. Fine-tuning the structure of glycosaminoglycans in living cells using xylosides. Glycobiology. 2018;28(7):499–511.

23. Weinstein T, Evron Z, Trebicz-Geffen M, Aviv M, Robinson D, Kollander Y, et al. β-D-xylosides stimulate GAG synthesis in chondrocyte cultures due to elevation of the extracellular GAG domains, accompanied by the depletion of the intra-pericellular GAG pools, with alterations in the GAG profiles. Connect Tissue Res. 20111207 ed2012. p. 169–79.

24. Hamati HF, Britton EL, Carey DJ. Inhibition of proteoglycan synthesis alters extracellular matrix deposition, proliferation, and cytoskeletal organization of rat aortic smooth muscle cells in culture. J Cell Biol. 1989;108(6):2495–505.

25. Carey DJ. Biological functions of proteoglycans: use of specific inhibitors of proteoglycan synthesis. Mol Cell Biochem. 1991;104(1-2):21–8.

26. Adamczyk P, Zenkert C, Balasubramanian PG, Yamada S, Murakoshi S, Sugahara K, et al. A non-sulfated chondroitin stabilizes membrane tubulation in cnidarian organelles. J Biol Chem. 2010;285(33):25613–23.

27. Faassen AE, Schrager JA, Klein DJ, Oegema TR, Couchman JR, McCarthy JB. A cell surface chondroitin sulfate proteoglycan, immunologically related to CD44, is involved in type I collagen-mediated melanoma cell motility and invasion. J Cell Biol. 1992;116(2):521–31.

28. Sutton A, Friand V, Brule-Donneger S, Chaigneau T, Ziol M, Sainte-Catherine O, et al. Stromal cell-derived factor-1/chemokine (C-X-C motif) ligand 12 stimulates human hepatoma cell growth, migration, and invasion. Mol Cancer Res. 2007;5(1):21–33.

29. Raman K, Kuberan B. Click-xylosides mitigate glioma cell invasion in vitro. Mol Biosyst. 2010;6(10):1800–2.

30. Mani K, Havsmark B, Persson S, Kaneda Y, Yamamoto H, Sakurai K, et al. Heparan/chondroitin/dermatan sulfate primer 2-(6-hydroxynaphthyl)-O-beta-D-xylopyranoside preferentially inhibits growth of transformed cells. Cancer Res. 1998;58(6):1099–104.

31. Lander C, Zhang H, Hockfield S. Neurons produce a neuronal cell surface-associated chondroitin sulfate proteoglycan. J Neurosci. 1998;18(1):174–83.

32. Pearson CS, Mencio CP, Barber AC, Martin KR, Geller HM. Identification of a critical sulfation in chondroitin that inhibits axonal regeneration. Elife. 2018;7.

33. Saied-Santiago K, Bülow HE. Diverse roles for glycosaminoglycans in neural patterning. Dev Dyn. 2018;247(1):54–74.

34. Kubota Y, Morita T, Kusakabe M, Sakakura T, Ito K. Spatial and temporal changes in chondroitin sulfate distribution in the sclerotome play an essential role in the formation of migration patterns of mouse neural crest cells. Dev Dyn. 1999;214(1):55–65.

35. Lafont F, Rouget M, Triller A, Prochiantz A, Rousselet A. In vitro control of neuronal polarity by glycosaminoglycans. Development. 1992;114:17–29.

36. Mutalik SP, Gupton SL. Glycosylation in Axonal Guidance. Int J Mol Sci. 2021;22(10).

37. Kantor DB, Chivatakarn O, Peer KL, Oster SF, Inatani M, Hansen MJ, et al. Semaphorin 5A is a bifunctional axon guidance cue regulated by heparan and chondroitin sulfate proteoglycans. Neuron. 2004;44(6):961–75.

38. Sasaki K, Komori R, Taniguchi M, Shimaoka A, Midori S, Yamamoto M, et al. PGSE is a novel enhancer regulating the proteoglycan pathway of the mammalian golgi stress response. Cell Struct Funct. 2019;44(1):1–19.

